# Highly Conserved Core Residues Define Old-World Alphaviruses and Trace Early Evolutionary Divergence

**DOI:** 10.1101/2025.09.25.678619

**Authors:** Wendy Carolina Piña-Ruiz, Luis Rubén Jaime-Rocha, Andrea Castorena-Robles, Roger Castells-Graells, Roya Zandi, Juan Carlos Muñoz-Escalante, Mauricio Comas-Garcia

**Affiliations:** High-Resolution Section, Research Center for Health Sciences and Biomedicine, Autonomous University of San Luis Potosi, Mexico; Department of Sciences, Autonomous University of San Luis Potosi, Mexico; Life Sciences Graduate Program, Department of Sciences, Autonomous University of San Luis Potosi, Mexico; Structural Biology Programme, Spanish National Cancer Research Centre (CNIO), Madrid, Spain; Department of Physics, University of California, Riverside; Bioinformatic Unit, Research Center for Health Sciences and Biomedicine, Autonomous University of San Luis Potosi, Mexico; Infectious Diseases Unit, Research Center for Health Sciences and Biomedicine, Autonomous University of San Luis Potosi, Mexico; Translational and Molecular Medicine Section, Research Center for Health Sciences and Biomedicine, Autonomous University of San Luis Potosi, Mexico

**Author notes:** To whom send any correspondence and. These authors contributed equally (Wendy Carolina Piña-Ruiz, Luis Rubén Jaime-Rocha).

**Keywords:** Alphaviruses, core evolution, capsid protein-capsid protein interactions, viral attenuation

## Abstract

Alphaviruses are positive-sense, single-stranded RNA viruses that assemble into striking double-icosahedral particles. During budding, their nucleocapsid core forms in the cytoplasm and adopts a T=4 icosahedral symmetry, a hallmark of Alphaviruses. Here, we combine structural and phylogenetic analyses to identify the amino acids most likely to govern capsomer formation (pentamers and hexamers) and core organization. We find that these residues are highly conserved in present-day Old-World alphaviruses but are divergent in New-World lineages. This suggests that the common ancestor of both groups likely assembled cores using the same molecular interactions seen in present-day Old-World viruses. We propose that early divergence in these interactions weakened core assembly efficiency, potentially contributing to the attenuation observed in encephalitic New-World alphaviruses. This attenuation may reflect an adaptive trade-off, in which reduced assembly efficiency lowers viral replication and virulence, supporting long-term persistence in enzootic cycles. Revealing how specific residues control capsid architecture and tracing their evolutionary history, this study provides key insights into alphavirus assembly mechanisms, opening new avenues for antiviral strategies and rational vaccine design.

## Introduction

Alphaviruses belong to the *Togaviridae* family, have a positive-sense single-stranded RNA genome of around 11-12 kilobases, and two open reading frames (ORFs) that code for two polyproteins (1). The ORF1 and ORF2 code the non-structural and structural proteins, respectively. Alphaviruses infect vertebrates and are transmitted by invertebrate vectors (2). These viruses have a double-layered virion consisting of two T = 4 icosahedral layers separated by a lipid membrane; the outer layer is composed of 80 trimeric spikes, each trimer composed of heterodimers of the glycoproteins E1 and E2, and the inner layer or core is made of 240 copies of the capsid protein (3). These spikes are not all placed at the same position in the icosahedron; 20 and 60 spikes at the 3- and quasi-3-fold symmetry axes, respectively. The core contains the full-length viral genome and assembles into an amorphous shell in the cytoplasm (4–7). At the plasma membrane, the core interacts with the heterodimer E1/E2, leading to the assembly and budding of the viral particle (4). Without the core, E1 and E2 bud as amorphous particles (8).

The Alphavirus virion has a unique architecture; the viral particle consists of two concentric T = 4 icosahedral layers separated by a lipid bilayer. Both layers interact through hydrophobic amino acids in the cytoplasmic domain of E2 and the capsid protein (CP) (3). These virions have structural features unique for an icosahedral viral particle(9–15): i) there are large “holes” at the center of the pentamer and hexamers (capsomers) of the core, ii) the capsomers do not establish large contact surface areas between them, and iii) the torsion angles and distances between capsomers can significantly differ between viral species.

Alphaviruses can be grouped by antigenic complexes (*e.g.,* Semliki Forest and Western equine encephalitis complexes) (16), by the location where they are endemic or first isolated (*i.e.,* Old- and New-World Alphaviruses) (17), or by some characteristic of the host (*e.g.,* if they infect aquatic animals or have no known vertebrate hosts) (18). Most Alphaviruses are classified as either Old- or New-World; the latter is restricted only to the American continent, and the former to the rest of the world. On the one hand, Old-World Alphaviruses mostly cause a febrile and arthrogenic dengue-like illness with a low mortality rate (19). On the other hand, New-World Alphaviruses affect the central nervous system, resulting in an encephalitic disease that, in some cases, has high mortality rates (up to 90% depending on the virus and the age of the patient) (20). Forrester and coworkers proposed that Alphaviruses originated from a marine host that jumped to terrestrial hosts, with birds being key to their global spread (18). Interestingly, some Alphaviruses are endemic to the New World (*i.e.,* Mayaro and Una viruses) but are phylogenetically part of the Old-World Alphaviruses. The opposite is true for the Sindbis virus, which is endemic to the Old World but is genetically related to New-World Alphaviruses (17, 18).

We first focused on identifying some structural differences in the cores of Alphaviruses using known structures to determine putative interactions that regulate CP-CP and capsomer-capsomer interactions. Then, we predicted the structure of the CP for all Alphaviruses and performed bioinformatic analysis of the consensus and representative amino acid sequences of the CPs. We found at least eight highly conserved amino acids in Old-World Alphaviruses, which are likely to mediate core assembly through electrostatic interactions (salt bridges). These amino acids are not conserved in other Alphaviruses. Finally, through a series of evolutionary analyses, we found that the core of the common ancestor most likely had almost the same electrostatic interactions as in Old-World Alphaviruses. These results suggest that a strong selective pressure resulted in the divergence of Old- and New-World Alphaviruses and altered the interactions of the core’s CP.

## Materials and Methods

### Viral sequences for the prediction of the capsid protein structure

The CP sequences for 42 *Alphavirus* species were obtained from the NCBI Taxonomy browser (Viruses; *Riboviria, Orthonovirae; Kitrinoviricota; Alsuviricetes; Martellivirales; Togaviridae*). For the E2 protein sequences, only 29 species had sufficient data to obtain representative sequences. The consensus sequences were generated using BioEdit v7.2.5.

### Structural data and structure modeling

The cryo-electron microscopy (cryo-EM) structures Sindbis (SINV) (9), Getah (GETV) (10), Chikungunya (CHIKV) (11), Mayaro (MAYV) (12), Western equine encephalitis (WEEV) (13), Venezuelan equine encephalitis (VEEV) (14), Eastern equine encephalitis (EEEV) (15), Barmah forest (BFV) (22), and Semliki forest (SFV) (23) viruses were obtained from the RCSB PDB website (with PDB IDs 3J0F, 7FD2, 3J2W, 7KO8, 8DEC, 7SFV, 6XO4, 2YEW, and 1DYl, respectively). The experimental resolution of these structures is 7.0, 2.8, 5.0, 4.4, 4.7, 4.0, 4.2, 5.0, and 9.0 Å, respectively. Models for the structures of the capsid protein and E2 were generated using AlphaFold2 (24); some sequences were randomly selected to be predicted using AlphaFold3 (25), and we obtained the same results in all cases as with AlphaFold2. The structures were analyzed and visualized using ChimeraX (26). All CP and E2 structures were compared to CHIKV (PDB ID: 3J2W) (11); in cases where multiple structures were obtained (*i.e.,* when using AlphaFold3), we reported the structure with the lowest RMSD when overlapped to CHIKV PDB ID 3J2W. All distances between side chains were calculated by measuring the centroid-to-centroid distances by generating spheres with 2 Å radius.

### Alphavirus sequences datasets curation for phylogenetic analysis

The complete nucleotide sequences corresponding to the 32 *Alphavirus* species were downloaded from GenBank using their taxonomy IDs. We used a different number of viruses for the structural prediction because the number of available sequences for some was not ideal for phylogenetic analyses. Sequences for each species were individually aligned using MAFFT v7.450 (27). The respective annotated RefSeq sequence for each species was used to identify the core coding sequence (CDS) region and to trim the multiple sequence alignments accordingly using BioEdit v7.2.5. To improve the quality of subsequent analyses, sequences were removed if they lacked more than 5% of the total length of the core CDS, contained more than 5% ambiguous nucleotides, or exhibited atypical indels not present in other sequences from the same species.

Consensus and representative sequences for each *Alphavirus* species were generated using CD-HIT-EST at a 90% similarity threshold. These representative core CDS sequences for each species were included in a single dataset and aligned using MEGA v10.2.6 with the MUSCLE algorithm (21). Amino acid sequences were inferred from the aligned nucleotide sequences, and manual curation of the Multiple Sequence Alignment (MSA) was performed using BioEdit v7.2.5.

### Phylogenetic analysis and reconstruction of ancestral sequences

Phylogenetic inference was conducted in MEGA v10.2.6 using the best-fitting model, and tree robustness was assessed with 1,000 bootstrap iterations. The most probable ancestral sequences for each internal node of the resulting phylogenetic tree were reconstructed using the ExtAncSeqMEGA extension.

### Selective pressure, Entropy, and Recombination analysis

To assess potential recombination events within *Alphavirus* CP coding sequences, alignments were analyzed using RDP v5.66, using RDP, GENECONV, BootScan, MaxChi, Chimaera, SiScan, and 3Seq as assessment methods. Potential events supported by at least three different methods with statistically significant *p-*values (<0.05) were considered true recombination events. Selective pressure acting on codon sites was evaluated on either all *Alphavirus* or only Old-World Alphaviruses nucleotide MSA using the Datamonkey Adaptive Evolution server employing SLAC, FEL, FUBAR and MEME methods; sites were classified as being under pervasive or episodic positive selection if detected by MEME (*p*<0.1), FUBAR (posterior probability >0.9), FEL (*p*<0.1), or SLAC (*p*<0.1), respectively.

Site-wise variability (either all *Alphavirus* or only Old-World Alphaviruses) was further examined using Shannon entropy calculation performed in BioEdit v7.2.5 on the MSA. Entropy scores were used to identify highly variable positions across the core protein sequences, supporting the detection of sites associated with adaptive or host-specific changes.

## Results

### The interactions between capsid proteins in Old-World viruses are highly conserved

First, we analyzed the cryo-electron microscopy (cryo-EM) structures of the core of Sindbis (SINV) (9), Getah (GETV) (10), Chikungunya (CHIKV) (11), Mayaro (MAYV) (12), Western equine encephalitis (WEEV) (13), Venezuelan equine encephalitis (VEEV) (14), Eastern equine encephalitis (EEEV) (15), Barmah forest (BFV) (22), and Semliki forest (SFV) (23) viruses (see Figure 1). In Figure 1, the different colors within a structure represent the four proteins that constitute the core asymmetric unit. These cores have a T = 4 symmetry; they are composed of 240 capsid proteins arranged in 12 pentamers and 30 hexamers. Despite having conserved overall architecture, close inspection of the hexamers, pentamers, and the three- and quasi-threefold symmetry axes shows interesting species-specific features (*e.g,* the number of amino acids establishing inter- and intra-capsomere interactions, the distance between capsomers and interacting amino acids, and the torsion angle between them).

**Figure 1.**
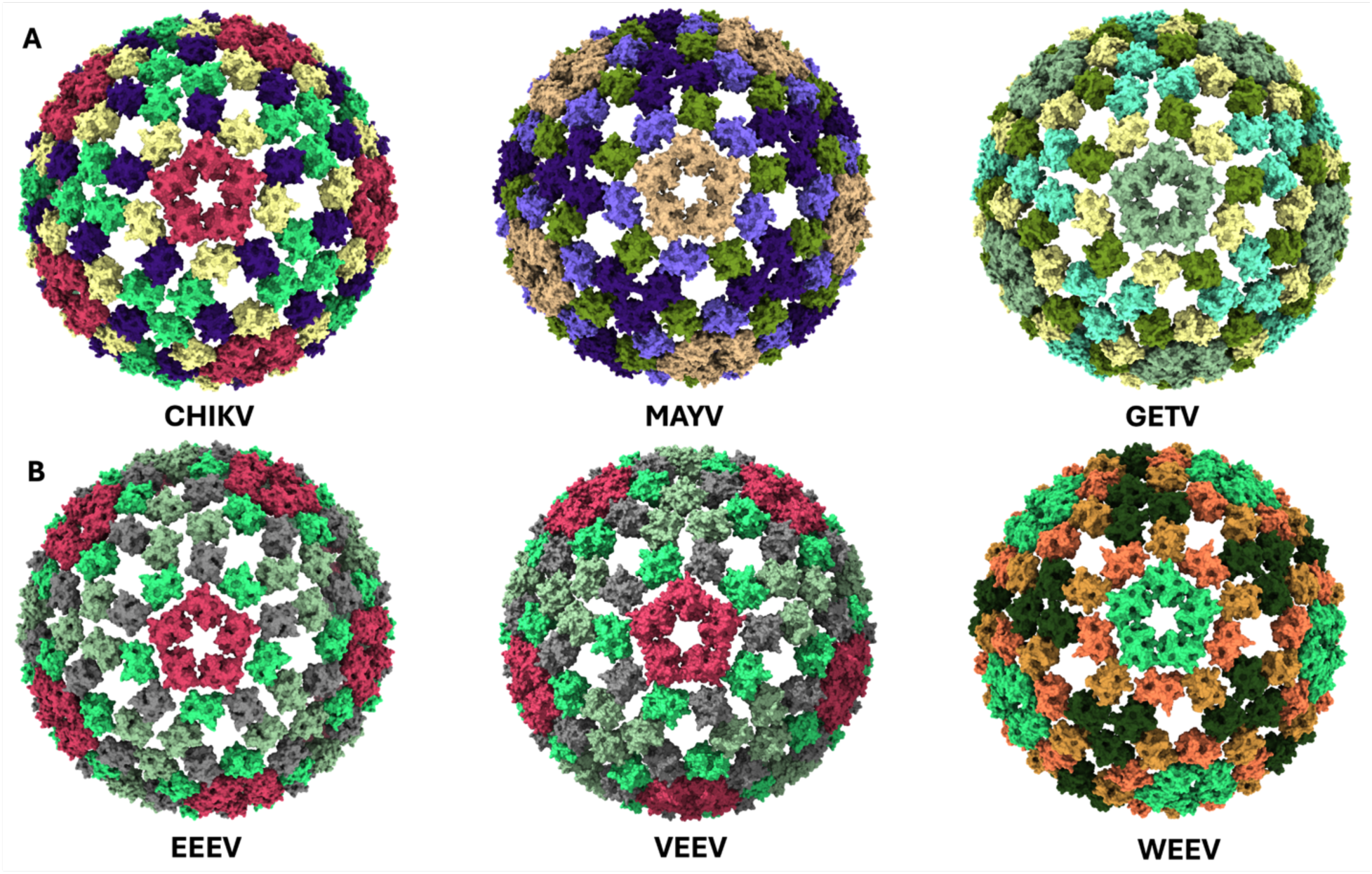
A) Comparison of the cryo-EM structures of the cores of CHIKV (PDB ID: 3J2W), MAYV (7KO8), and GETV (7FD2). B) Comparison of cryo-EM structures of the cores EEEV (6XO4), VEEV (7SFV), and WEEV (8DEC). Each structure has four diSerent colors, representing that in a T = 4 icosahedron, the capsid protein (CP) in the asymmetric unit has four diSerent chemical environments.

Figure 2A shows no significant differences in the pentamers of these three Old-World Alphaviruses (*i.e.,* CHIKV, MAYV, and GETV) compared to those in the three New-World Alphaviruses in Figure 2B; in EEEV, the capsid proteins (CP) in the pentamer establish fewer inter-subunit contacts than in VEEV and WEEV (see the asterisks). The structures of the VEEV and WEEV pentamers exhibit no noticeable differences compared to those of CHIKV, MAYV, and GETV. However, the quasi-threefold symmetry axis (q3) shows some changes (supplementary figure 1); the space between two adjacent hexamers and one pentamer is approximately 23 Å, 19 Å, and 25 Å wide for CHIKV, MAYV, and GETV, respectively. These distances were calculated by generating 2 Å centroids for the amino acids K122, K119, and K129 in CHIKV, MAYV, and GETV, respectively. Then we measured the centroid-to-centroid distance between these amino acids in the three proteins forming this symmetry axis. Amino acids Q121, K125, and Q119 were used for EEEV, VEEV, and WEEV, respectively (in the following sections we will explain why we used those amino acids as references). Interestingly, while the general features of the VEEV pentamer are like those of the Old-World Alphaviruses, the space at the q3 is much smaller (≈ 13 Å) than for the other five viruses in this figure (≥ 15 Å) (see supplementary figure 1). The three-fold symmetry axis (*i.e.,* between three hexamers) shows a pattern (data not shown).

**Figure 2.**
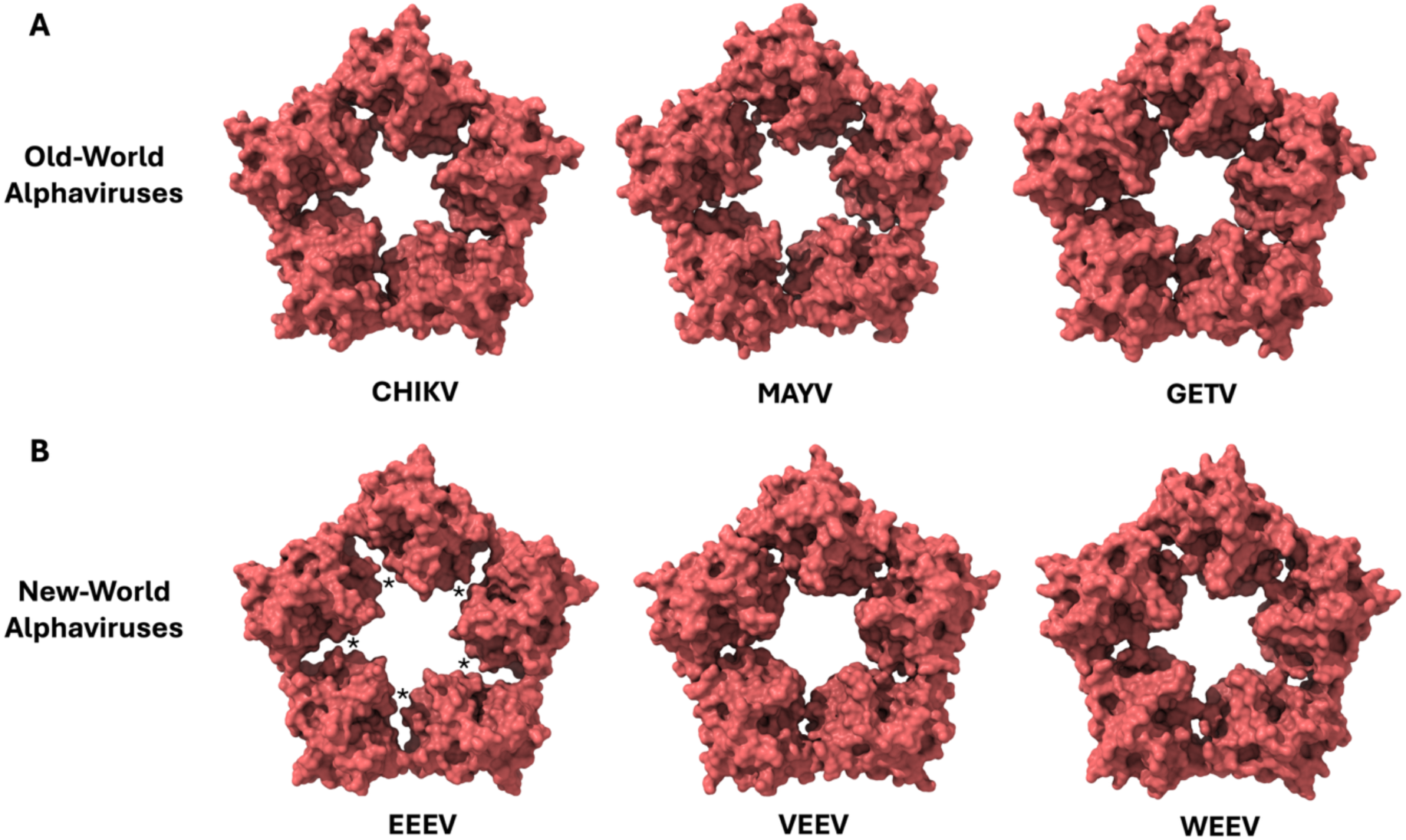
Comparison of the structure of the pentamers with the structures in Figure 1. The proteins in the pentamer have the same color because they share the same chemical environment, unlike the proteins in the hexamer, which have three distinct chemical environments. A) The structures of the pentamers of these three Old-World Alphaviruses are very similar. B) The structure of the EEEV pentamer has fewer contact angles (asterisks) than the other New-World Alphaviruses. The contacts between CPs in VEEV and WEEV are similar to those of CHIKV, MAYV, and GETV.

These results suggest that the physicochemical properties of the amino acids, which should control capsomer assembly and capsomer-capsomer interactions, may be similar for these three Old-World viruses and vary in the New-World Alphaviruses. To explore this idea, we focused on the CP contact surfaces, using the CHIKV core cryo-EM structure (PDB ID 3J2W) as a reference. The assembly of Alphaviruses occurs at neutral pH at the plasma membrane, while disassembly occurs in the endosome under acidic conditions (3). Therefore, we focused on pH-dependent interactions in multiple CP-CP interfaces using CHIKV as our reference system.

Figure 3A suggests electrostatic interactions (salt bridges) between CHIKV CP K172 and E234 (outer), and K177 and E184 (inner). It should be noted that the resolution of these structures does not permit precise localization of side-chain positions. Nonetheless, the distances of these amino acids and their chemical environment suggest electrostatic interactions between the CPs in the pentamers. These putative interactions are also present in the hexamers (data not shown); however, because of the asymmetry of the hexamer, the distances between K177 and E184 vary between adjacent CPs and are greater than in the pentamers (data not shown). Figure 3B shows the three-fold symmetry (i3) axis of CHIKV, suggesting that the capsomer-capsomer interactions depend on two sets of electrostatic interactions: E120/K151 and K122/D148. The position of these amino acids resembles a dipole-dipole interaction; thus, the strength of these interactions should depend on the distances and angles between capsomers. To further understand the structures shown in Figure 2, we overlapped the structure of the monomeric CHIKV CP (Figure 3C) with those of Alphaviruses, MAYV and EEEV, with a known cryo-EM structure. Figures 3D and 3E show the overlap of CHIKV CP with MAYV and EEEV CPs, respectively. On the one hand, Figure 3D shows that MAYV CP maintains perfect charge conservation and identity at the positions of the eight amino acids of interest. The only difference is that CHIKV CP K151 has an arginine in MAYV (R148). On the other hand, Figure 3E shows key differences in the EEEV CP; the amino acids that should be in the same position as in CHIKV CP K122, K151, D148, K177, and E184 are instead Q, Q, A, L, and P, respectively. The overlap of the structures of GETV, SFV, and BFV with CHIKV shows that the amino acids in these eight positions are 100% conserved in charge, while the degree of conservation for SINV, WEEV, and VEEV is much lower (data not shown).

**Figure 3.**
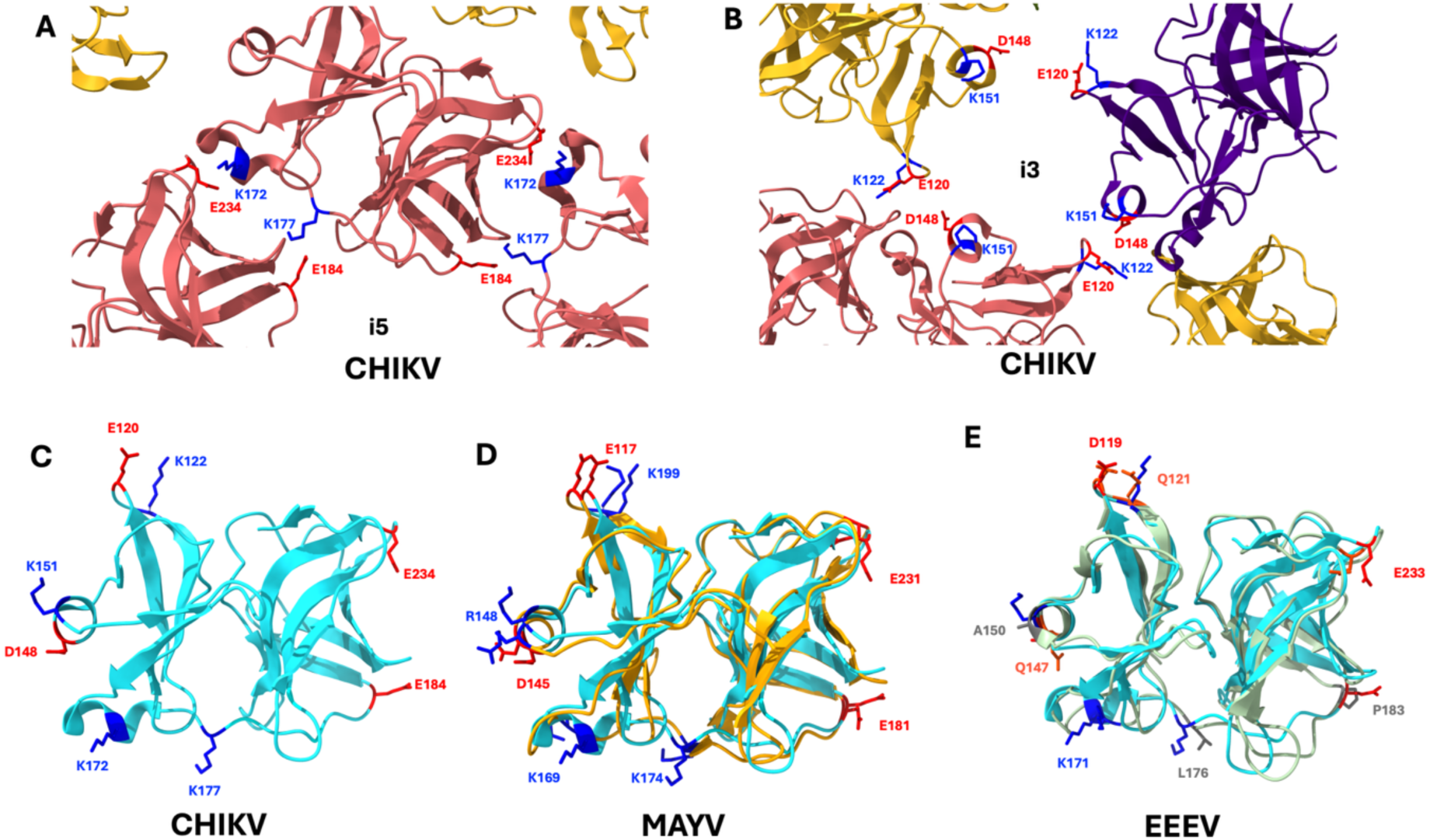
A) Putative electrostatic interactions at the surface between three proteins in the CHIKV pentamer, between E184 and E234 of one protein and K172 and K177 of the other protein. B) The interactions at the quasi-threefold symmetry axis resemble dipole-dipole interactions between the pair of amino acids E120/K122 and D148/K151. C) CHIKV CP (PDB ID: 3J2W) monomer displaying E120, K122, D148, K151, K172, K177, E184, E234. D) Overlap of CHIKV CP with MAYV CP (PDB ID: 7KO8), where, although the numbering and chemical identity of some amino acids differ, the charges are conserved in all amino acids. E) Overlap of CHIKV CP with EEEV CP (PDB ID: 8DEC). This overlap indicates that some amino acids are not conserved in EEEV, suggesting that the intra- and inter-capsomere interactions in EEEV diSer significantly from those in MAYV and CHIVK.

Figure 4 shows a closer look at the interfaces in a pentamer; Figure 4A (top row) contains only Old-World Alphaviruses, which exhibit the same salt bridges as CHIKV. Figure 4B shows the equivalent interface for three New-World Alphaviruses; the amino acids in positions equivalent to CHIKV CP K177 and E184 (inner interactions) are very different between EEEV, VEEV, and WEEV. Nonetheless, the charge of the amino acids equivalent to CHIKV CP K172 and E234 (outer interactions) is the same in all cases. The “outer interactions” in the pentamer seem to be conserved in Old- and New-World Alphaviruses, while the “inner interactions” are only conserved in Old-World Alphaviruses. Interestingly, while there are large differences in the CP conformation and the CP-CP surfaces in the pentamers of Old-World Alphaviruses, the “inner” and “outer” electrostatic interactions seem to be the same. In the New-World Alphaviruses, the CP conformation, the CP-CP surfaces, and the putative interacting amino acids that are equivalent in position to those in Old-World Alphaviruses are not conserved.

**Figure 4.**
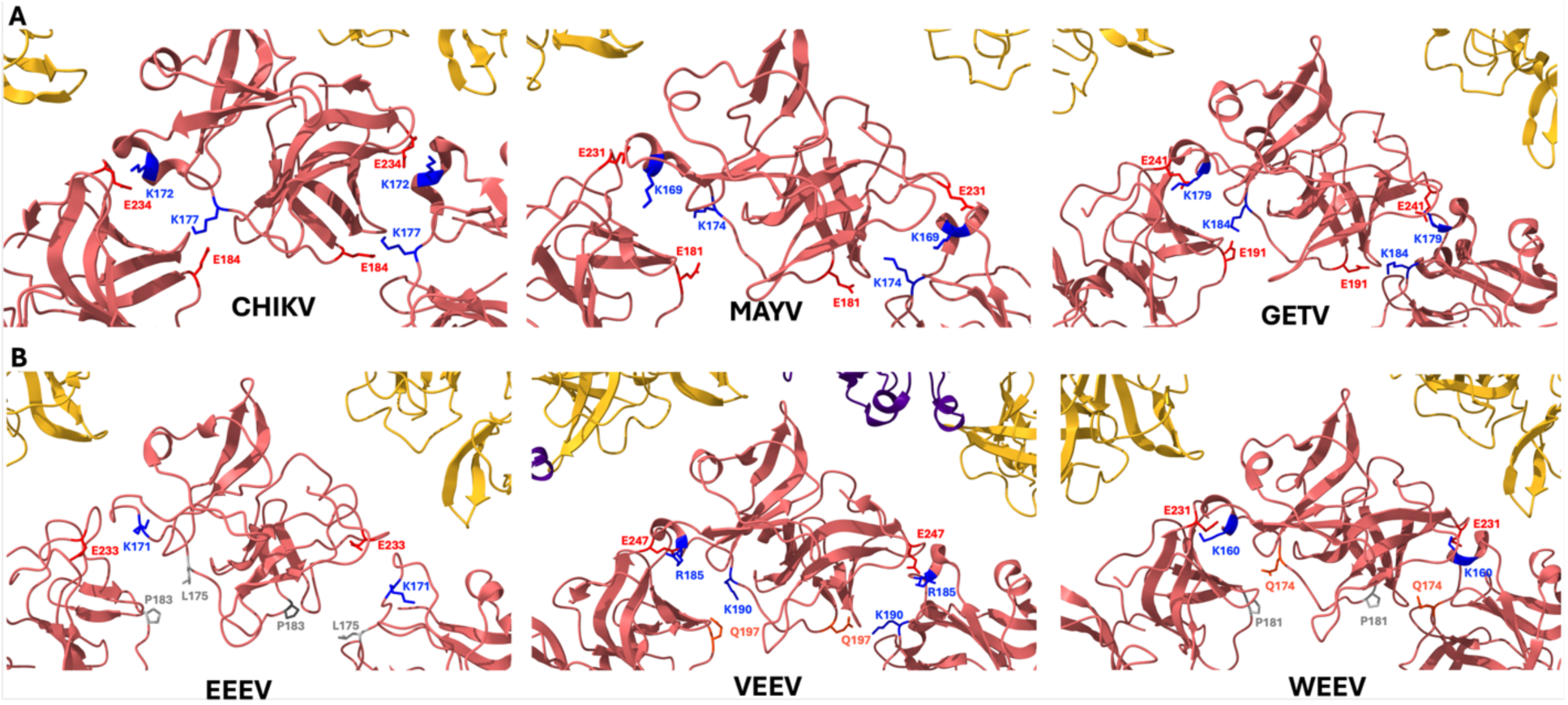
Comparison of the interfaces between the three proteins at i5 (pentamer). A) The “outer interactions” for CHIKV are between K172 and E234, and the “inner interactions” are between K177 and E184. It also shows that the possible interactions between amino acids at equivalent positions at the same interfaces for MAYV and GETV are the same as in CHIKV. B) The “outer interactions” for EEEV, VEEV, and WEEV are the same as in the Old-World Alphaviruses from the top row. However, the “inner interactions” are diSerent. The inner interactions for EEEV involve a P and L, which could explain the large holes observed in Figure 2. In VEEV, the amino acids at the “inner position” are Gln and Lys, while for WEEV, they are P and Q.

To better understand the interactions between capsomers (Figure 3B), we analyzed the i3 axis (see Figure 5). The leftmost panel in Figure 5A shows the centroid-to-centroid distance (2-Å centroid) and torsion angle between three CHIKV CP K122 amino acids. The letters in parentheses indicate the chains being compared in the Cryo-EM structure. The distances between equivalent amino acids are asymmetric, although the exact positions of the side chains are unknown. This analysis was also conducted for MAYV, GETV, EEEV, VEEV, and WEEV. The interactions in CHIKV, MAYV, and GETV (Figure 5A) are the same in these viruses, and the distances and torsion angles between equivalent amino acids do not differ between these viruses. Figure 5B shows that at the i3, the interactions in EEEV and WEEV are between polar uncharged amino acids (Q and N) instead of charged amino acids.

**Figure 5.**
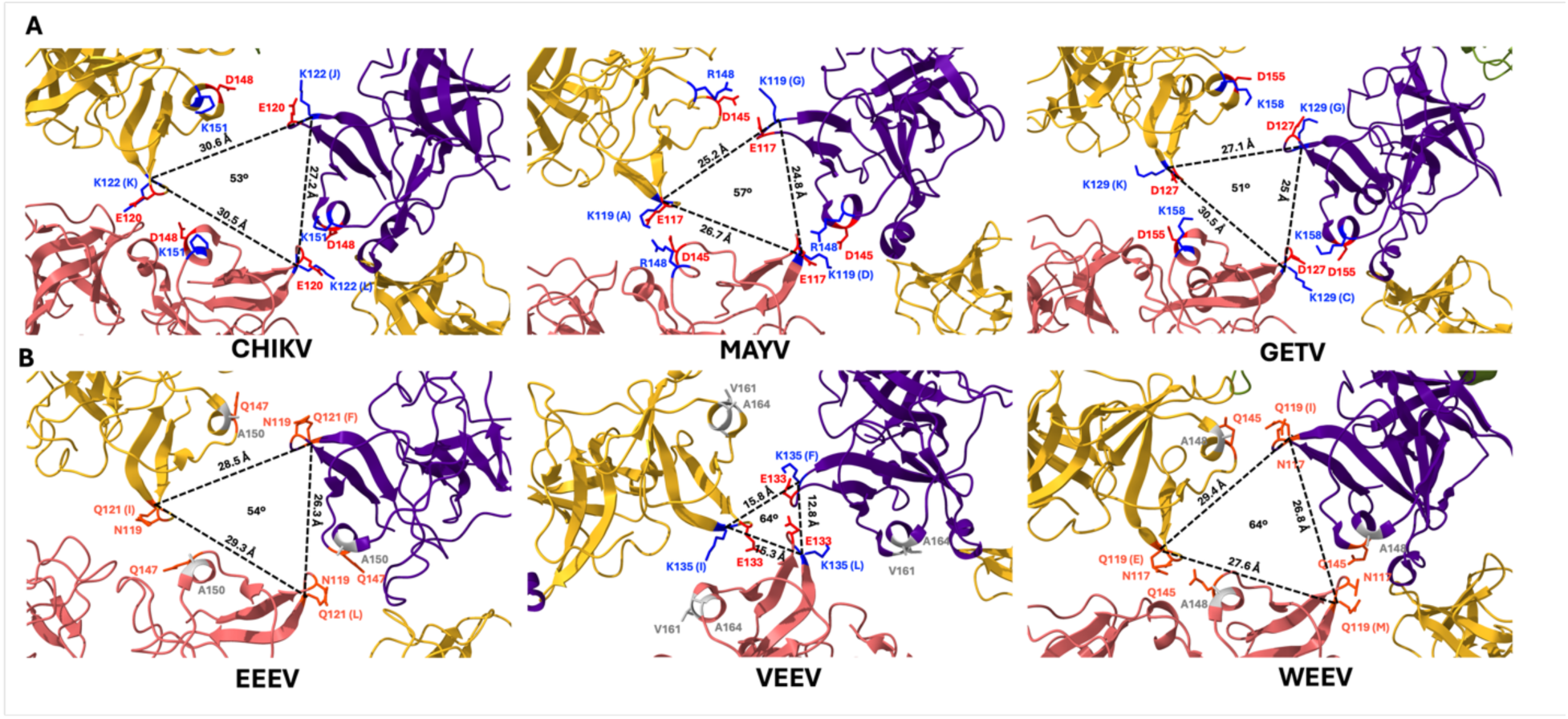
A) The leftmost panel shows the geometry of the i3 axis for CHIVK, MAYV, and GETV, which was determined by measuring the centroid-to-centroid distance (2-Å centroid) and torsion angle between K122 of the three proteins at this axis. B) The distances between these three viruses and EEEV and WEEV, as well as those in the top row, are very similar. Only VEEV has a distance that is much smaller than that of the other viruses. Also, the distances for EEEV and WEEV were calculated by using Q121 and Q119, respectively, which are in equivalent positions to CHIKV CP K122. Interestingly, in VEEV, K134 and E133 do not have a pair of charged amino acids, as seen in the top row, that could help establish a dipole-dipole-like interaction. The torsion angles between three equivalent amino acids are indicated at the center of the i3 axes. The torsion angles for CHIKV, MAYV, and EEEV are very similar; however, VEEV and WEEV have larger torsion angles than the other four viruses.

Additionally, the middle panel in Figure 5B shows that in VEEV, a cluster of charged amino acids (K135 and E135 from three capsid proteins) does not appear to have a corresponding pair of amino acids that could establish a dipole-dipole-like interaction, as observed in CHIKV, MAYV, and GETV (see Figure 5A). The distances between Q121 and Q119 in EEEV and WEEV are similar to the basic amino acids in the i3 of CHIKV, MAYV, and GETV. The only significant difference in WEEV is that the torsion angle of the amino acid is greater than that of Old-World Alphaviruses. The interactions in VEEV are very different from those of the other viruses in this Figure; for example, the charged amino acids are about half the distance as in CHIKV (≈ 12.8-15.8 Å vs ≈ 27.2-30.6 Å, respectively). There is no pair of amino acids with an opposite charge that could help to establish a dipole-dipole-like interaction. The torsion angles between these amino acids are indicated in the center of the i3 site; these angles are very similar for these three Old-World Alphaviruses and EEEV (between 51-57 °). However, EEEV and WEEV have different torsion angles, albeit having similar distances and amino acids at the sites of interest. This suggests that other amino acids contribute to the CP-CP in these interfaces.

### The electrostatic interactions between capsid proteins are a unique feature of Old-World viruses

The results in the previous section were limited to Alphaviruses with known Cryo-EM structures; however, we wanted to identify these interactions for all Alphaviruses with known CP sequences. First, we generated a consensus sequence for the CP of all Alphaviruses (including those in the previous section) and predicted the structure of the monomeric capsid protein using Alphafold2 (24). We also used Alphafold3 (25) for a few randomly selected structures, and the results were consistent with those obtained with Alphafold2. The goal of this prediction was to identify amino acids for Alphaviruses without structural data that are in positions equivalent to those of interest in the monomeric form of the CHIKV CP and not to infer any data from a possible multimeric context. In fact, we only analyzed the predicted monomeric structures of the capsid proteins because Alphafold2 and 3 failed to assemble them into pentamers or hexamers. For this analysis, we used consensus sequences. Then, we overlapped the predicted structures with the CHIKV CP (PDB 3J2W) as was done in Figures 3D and 3E to identify the amino acids that are in equivalent positions to those of CHIKV CP, specifically E120, K122, D148, K151, K172, K177, E184, and E234 (see supplementary files). From here on, we will use this numbering to indicate the amino acids of reference, and CHIKV will be the virus to which we will compare all interactions. The results of these analyses are presented in Figures 6-9. The numbers in green indicate that the percentage of amino acid identity and/or charge conservation is greater than or equal to 80%.

**Figure 6.**
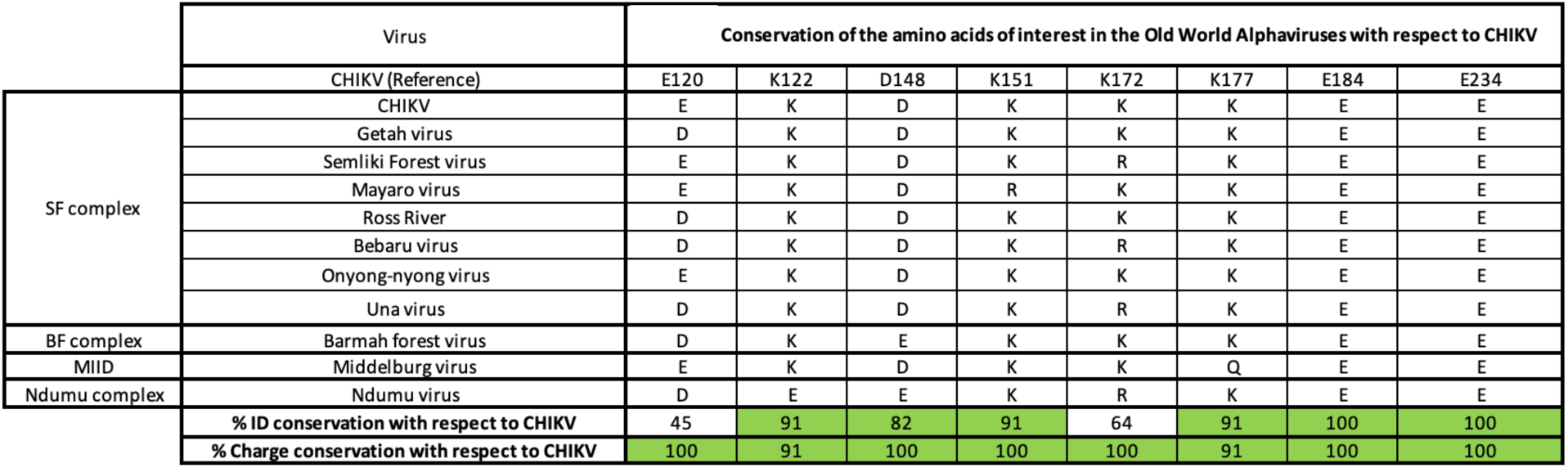
Comparison of the identity of amino acids in equivalent positions to CHIKV CP E120, K122, D148, K151, K177, E184, and E234 for Old-World Alphaviruses. These amino acids were identified by predicting the structure of the consensus CP sequence of all viruses using AlphaFold2. The green color indicates that the charge and/or identity of the amino acids are conserved at least 80%.

Additionally, we performed a multiple-sequence alignment (MUSCLE) of the consensus sequences (see Supplementary Figures 2 and 3). The comparison between the two methods revealed a few differences. The lavender, turquoise, and maroon amino acids in Figures 7-9 indicate that the positions of the amino acids predicted through Alphafold2 and MUSCLE differ. The lavender color indicates that the amino acid overlapping with the reference structures is due to an insertion. The amino acids in turquoise overlap at a -1 position in the aligned sequence, and the maroon color refers to amino acids that overlap with the reference CHIKV amino acids because of a deletion.

**Figure 7.**
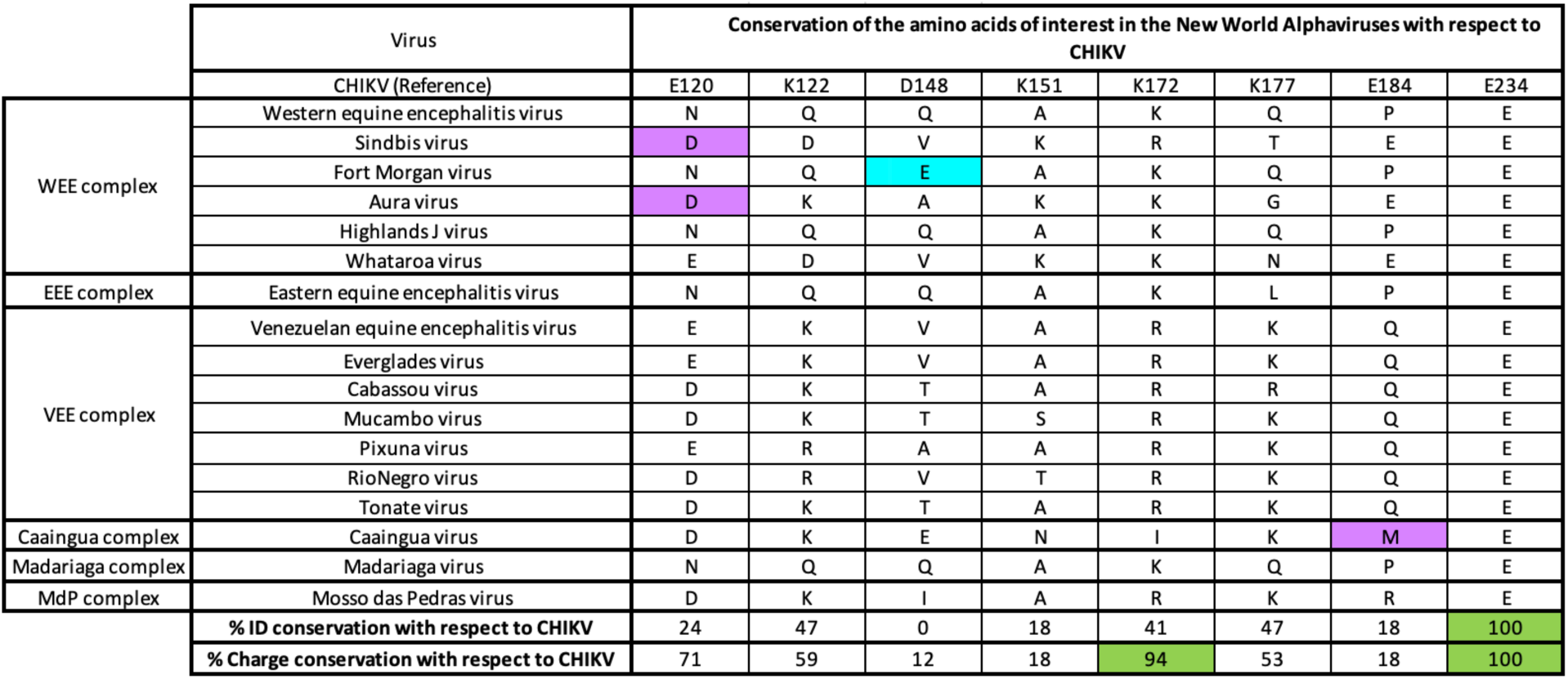
Comparison of the identity of amino acids in equivalent positions to CHIKV CP E120, K122, D148, K151, K177, E184, and E234 for New-World Alphaviruses. These amino acids were identified, as explained in Table 1. The lavender and turquoise colors indicate that these amino acids are at equivalent positions to those in CHIKV due to an insertion or a -1 shift, respectively. Only the positions equivalent to K172 and E234 are conserved at 94% and 100%, respectively.

Figure 6 shows almost 100% charge conservation of these amino acids in all Old-World Alphaviruses; the Ndumu and Middelburg viruses have a charge inversion in K122 and a Q in K177, respectively. Also, except for the positions equivalent to E120 and K172, the identities of the other six amino acids are conserved between 80% and 100%. These results suggest that for Old-World Alphaviruses, there is strong evolutionary pressure on the charge of amino acids that might control the intra- and inter-capsomer interactions and the length of the side chains of these amino acids. These findings are consistent with the results presented in the previous section.

In New-World Alphaviruses, except for the amino acids equivalent to K172/E234 (*i.e.,* the “outer interactions”), the charge is not conserved in the rest of the amino acids of interest (Figure 7). However, there seems to be some degree of conservation within antigenic complexes. In most cases, the charged amino acids are replaced by polar uncharged amino acids, except in the positions equivalent to D148 and K151, where there is a significant amount of hydrophobic side chains. Interestingly, E120 and K122 are conserved in the VEEV antigenic complex. In contrast, in the WEEV complex, E120 can be a polar uncharged, or acidic side chain, and K122 is either an acidic or a polar uncharged amino acid. In WEEV and VEEV, D148 is mostly substituted for Q or V. In the WEEV complex, K177 and E184 are Q and P, respectively, while in the VEEV complex, the charge of K177 is conserved, and E184 is almost always a Q. The identities of five of the eight amino acids of interest within the VEEV complex are more conserved than within viruses of the WEEV complex. These results suggest a common ancestor for the CP of Aura virus, SINV, and Whataroa virus and a distinct ancestor for the other viruses within this antigenic complex. Furthermore, based on the results of Table 2, it is likely that the CP of the Madariaga virus may have had a common ancestor with the cores of WEEV, Fort Morgan Virus, and Highland J virus.

The lack of conservation of intra- and inter-capsomeric interactions is more evident for Alphaviruses found in mosquitoes with no known vertebrate host that are not included in the Old-/New-World classification and on aquatic Alphaviruses (see Figures 8 and 9). Figure 8 shows that, on the one hand, like in New-World Alphaviruses, a hydrophobic side chain replaces D148. On the other hand, K151 is replaced by a polar uncharged amino acid, which is unique for this group. Interestingly, the charge conservation of the viruses in Table 3 is closer to that of Old-World than New-World Alphaviruses.

**Figure 8.**
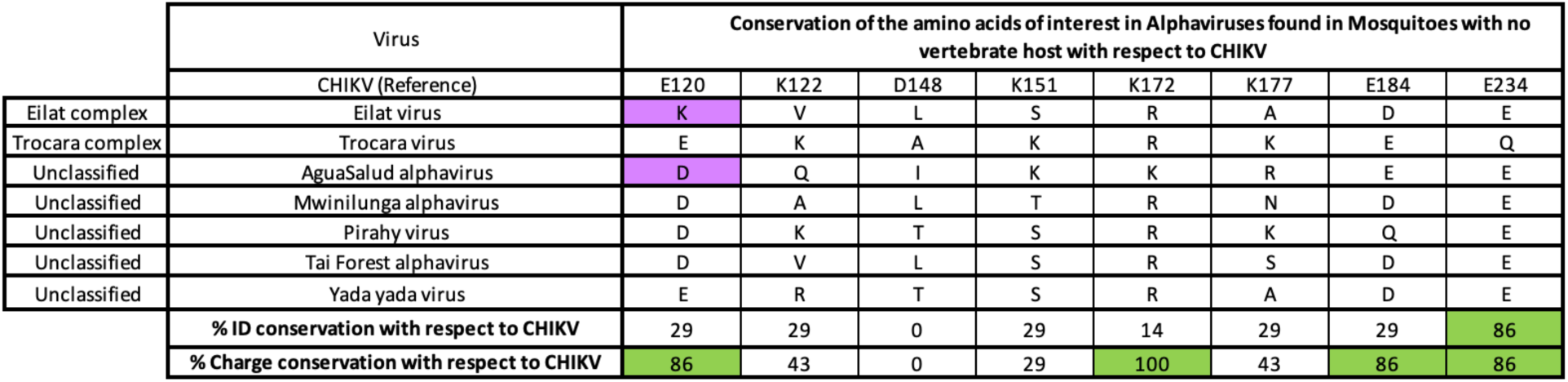
Comparison of the identity of amino acids in equivalent positions to CHIKV CP E120, K122, D148, K151, K177, E184, and E234 for Alphaviruses with no known host is not part of the Old-/New-World classifications. These amino acids were identified, as explained in Table 1. The lavender color indicates that those amino acids are at equivalent positions to CHIKV due to the insertion. Only the positions equivalent to K172 and E234 are conserved at 94% and 100%, respectively. In this group, positions K172 and E234 are conserved, as well as E120 and K172.

The degree of charge conservation of the amino acids of interest in aquatic Alphaviruses is low compared to CHIKV. However, in aquatic Alphaviruses that infect mammals, six of the eight amino acids of interest are conserved in these viruses (see shaded rows in Figure 9). While the number of viruses in this category is too small to make a conjecture, it could suggest that, at least based on the core and/or CPs, these two viruses might be related to Old-World Alphaviruses. In aquatic Alphaviruses that infect fish, there are two positions with charge inversion, K151 and K177. This charge inversion should result in intra- and inter-capsomeric interactions that differ from the rest of the Alphaviruses.

**Figure 9.**
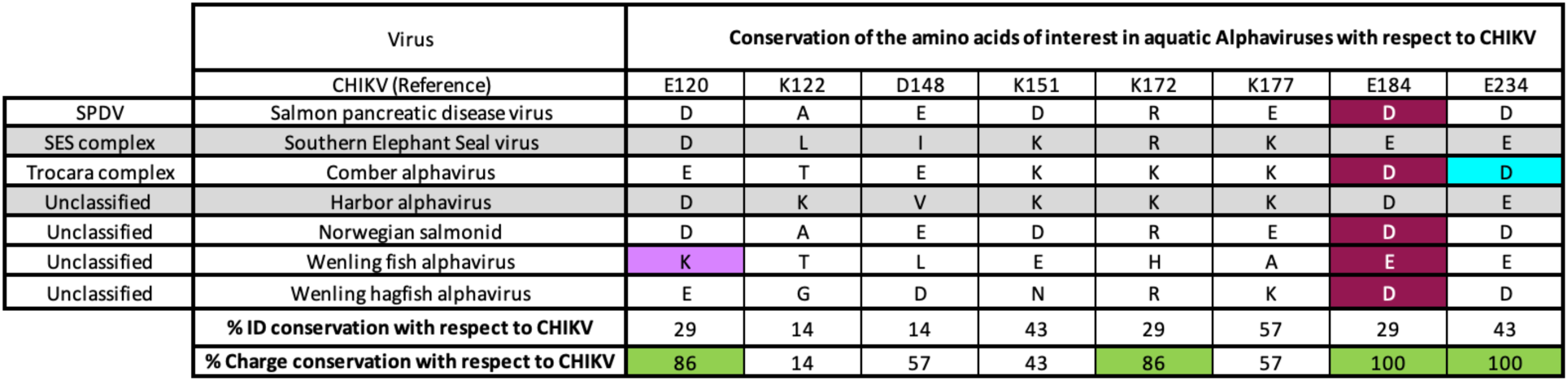
Comparison of the identity of amino acids in equivalent positions to CHIKV CP E120, K122, D148, K151, K177, E184, and E234 for aquatic Alphaviruses. These amino acids were identified, as explained in Table 1. The lavender, turquoise, and maroon colors indicate that those amino acids are at equivalent positions to CHIKV due to an insertion, a -1 shift, or a deletion, respectively. Positions K172, E234, E120, and K172 are conserved. The gray color indicates that those viruses infect mammals. The rest infect fish.

The data in Figures 6-9 show a wide range of interactions for non-Old-World Alphaviruses. Nonetheless, it is clear that the “outer intracapsomeric interactions” (E234/K172) are highly conserved in almost all Alphaviruses; only in the Trocara virus and the Wenling fish alphavirus, the amino acids in the equivalent positions to CHIKV CP E234 and K172 are a Q and an H, respectively. These results suggest that there is a strong evolutionary pressure on the salt bridge between CHIKV CP E234 and K172, and that this interaction has been conserved during evolution, possibly playing a crucial role in core assembly.

### The electrostatic interactions between capsid proteins at the core might be related to glycosylation patterns in E2

The results presented so far suggest that the organization and type of interactions in the core of Alphaviruses could have changed when Old- and New-World Alphaviruses diverged. But these are not the only changes. The glycosylation profiles of E2 and E1 for Old-World Alphaviruses differ from New-World ones (28). For example, CHIKV E2 has only one glycosylation site in N623. A bioinformatic and structural analysis identical to the one described in the previous section, aiming at finding an N-X-S/T motif at the same position as CHIKV E2 N263, shows that this glycosylation site is conserved in almost all Old-World Alphaviruses (see Figure 10A). Only the Bebaru virus appears to have an N-to-G substitution at this site. We could not find an N-X-T/S motif in or around the E2 region of the Bebaru virus that is structurally equivalent to the CHIKV motif. Figure 10B shows the Cryo-EM density of the E2 glycosylation site for MAYV, which has been proposed to act as a “molecular handshake” between trimers that stabilize the viral particle (12). It should be noted that analyses E2 and CP do not have the same number of species, as there are fewer available sequences in E2 than in CP. Therefore, the number of viruses in Figures 6-9 and Figures 10A, 10B, and 10C differs, and thus, this could limit our study.

**Figure 10.**
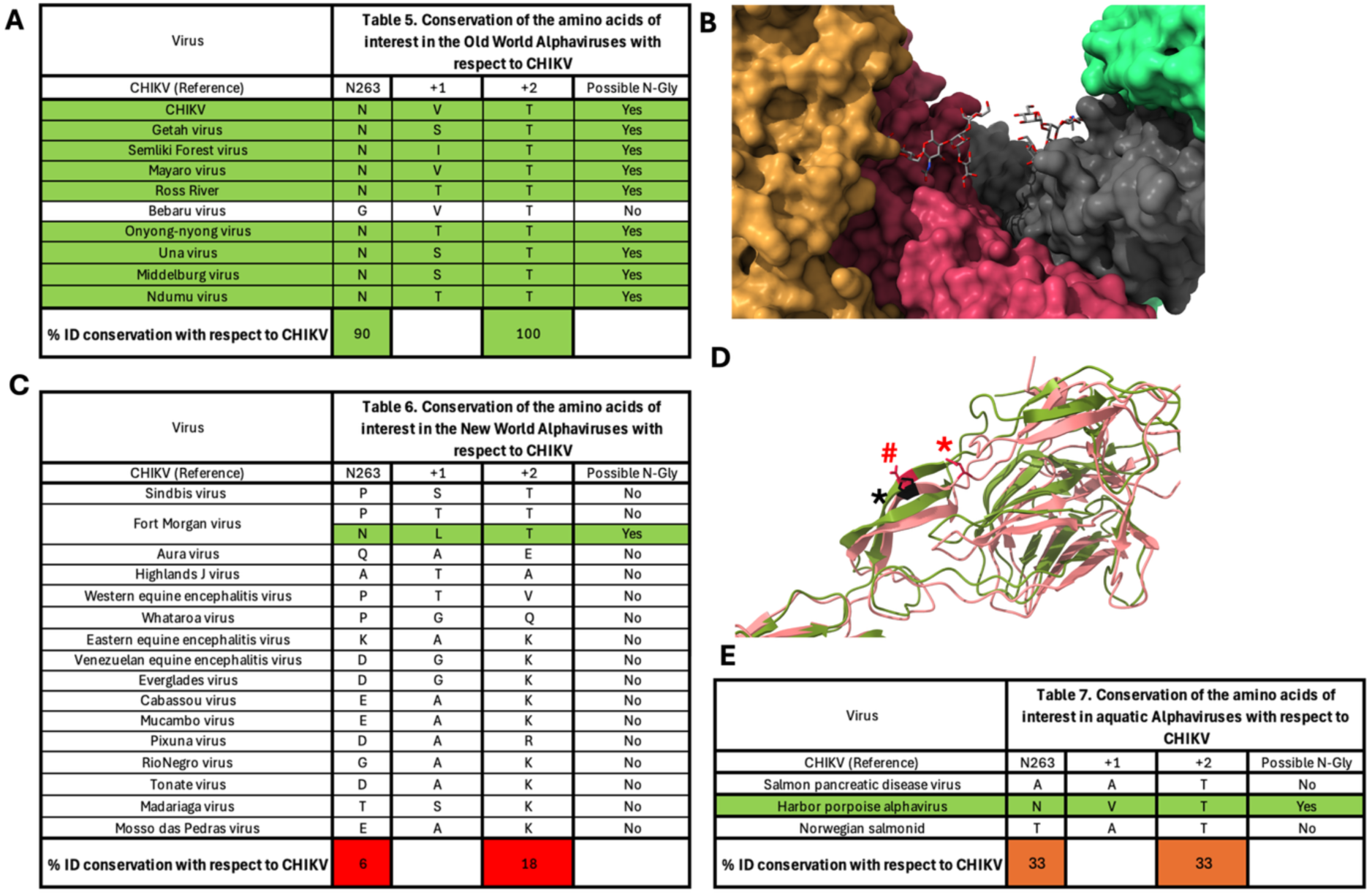
A) Except for the Bebaru virus, all other Old-World Alphaviruses in this table have a predicted N-glycosylation site at a position that is equivalent to CHIKV N263. B) Cryo-EM structure of MAYV virus (PDB ID: 7KO8) showing the “molecular handshake” between glycosylation sites. This interaction occurs between two trimers of heterodimers. C) New-World Alphaviruses do not have a predicted N-glycosylation at an equivalent position to CHIKV N263. The only exception is the Fort Morgan virus, which has a possible glycosylation site at N260, which is not in the same position as CHIKV E2 N263. D) Overlap of CHIKV E2 (green) with the predicted structure for the Fort Morgan virus E2 protein (salmon). CHIKV E2 N263 is in red and indicated by a red # and is in a β-sheet. The amino acid in the Fort Morgan virus E2 protein equivalent to CHIKV E2 N263 is in black and indicated by a black asterisk (Proline), while the Asp in an N-L-T motif is in a loop and is indicated with a red asterisk. If this amino acid were glycosylated, it is unlikely to be able to establish the same interactions as the glycosylation site shown in panel B. E) From the Alphaviruses that infect aquatic animals with available E2 sequences, only the Harbor Alphavirus has the same predicted E2 glycosylation site as CHIKVK. Interestingly, the core of this virus has seven of the eight interactions conserved in the CPs of Old-World Alphaviruses, as shown in Figures 4, 5, and 6 and Tables 1 and 4. This small correlation could suggest that the nature of the core and the interactions between trimers are related to each other.

Figure 10C shows no N-glycosylation motif on or around CHIKV E2 N263 in New-World Alphaviruses. This figure also suggests that the Fort Morgan virus may have an N-glycosylation site (N260) in a position close to CHIKV E2 (N263). However, E2 N260 in the Fort Morgan virus is not equivalent to CHIKV E2 N263. In the Old-World Alphaviruses, the glycosylation site equivalent to CHIKV N263 is in a β-sheet, while the Fort Morgan N260 (N-V-T motif) is in a loop before this β-sheet (Figure 10D). The position of this possible glycosylation site makes it unlikely to result in the same trimer-trimer interactions as the “molecular handshake” proposed for MAYV (12).

Finally, the data in Figure 10E suggest that Harbor porpoise alphavirus, which infects marine mammals, has the same glycosylation site as CHIKV. Interestingly, this marine Alphavirus has seven of the eight amino acids studied in the previous sections conserved in charge compared to CHIKV. Therefore, there may be a relationship between the identity of the amino acids that control assembly and the position of the glycosylation sites in E2. It is possible that in non-Old-World Alphavirus cores, the interactions between CPs are weak, and glycosylation in E2 N263 could introduce rigidity that inhibits the assembly of the icosahedral core during budding.

### The common ancestor of Old- and New-World Alphaviruses may have had intra- and inter-capsomer interactions like those of CHIKV

Phylogenetic inference analysis of Alphaviruses is usually done using the structural viral protein E1, E2, and TF, or a non-structural protein (17, 18, 29). However, using E1 and E2 provides information about the evolution of viruses in the context of their interactions with hosts. Therefore, using these proteins is extremely useful for understanding the evolution of a pathogen from a perspective that incorporates contributions from both the host and vectors. Nonetheless, we wanted to understand the evolution of the core and determine what kind of core was first: was the core with strong electrostatic interactions (Old-World Alphaviruses) first, and then the core with weak interactions appeared (New-World Alphaviruses), or vice versa? First, we generated representative viral sequences with sufficient CP data and performed phylogenetic inference using MEGA v10.2.6 and ExtAncSeqMEGA (see Methods and Materials).

Figure 11 shows the result of the phylogenetic analysis using CP. The structural analysis was conducted using consensus sequences; thus, we studied 42 viral species. However, to obtain a robust phylogenetic analysis, we used representative sequences, which limited us to 32 species. As expected from the results shown in Figures 6 and 7, there is a clear separation between Old- and New-World Alphaviruses; however, this analysis shows that the CP of Sindbis, Aura, and Whataroa viruses diverged separately from the rest of the New-World clades. This separation is similar but not identical to those reported using the structural proteins E1, E2, and TF (17, 18, 29). Figures 9 and 10D suggest that Harbor porpoise alphavirus did not jump directly from fish to a porpoise, but it appeared when the Old-World group separated from the Aura virus clade (blue clade in Figure 11). As shown by Weaver and Forrester (17, 18), Middelburg, Bebaru, Semliki, Getah, and Ross River viruses belong to a distinct clade, separate from CHIKV, O’nyong nyong, Una, and Barmah Forest viruses. However, our results differ in some respects. By looking only at the CP, the Una and Mayaro viruses are not part of the same clade as proposed based on E1, E2, and TF (17).

**Figure 11.**
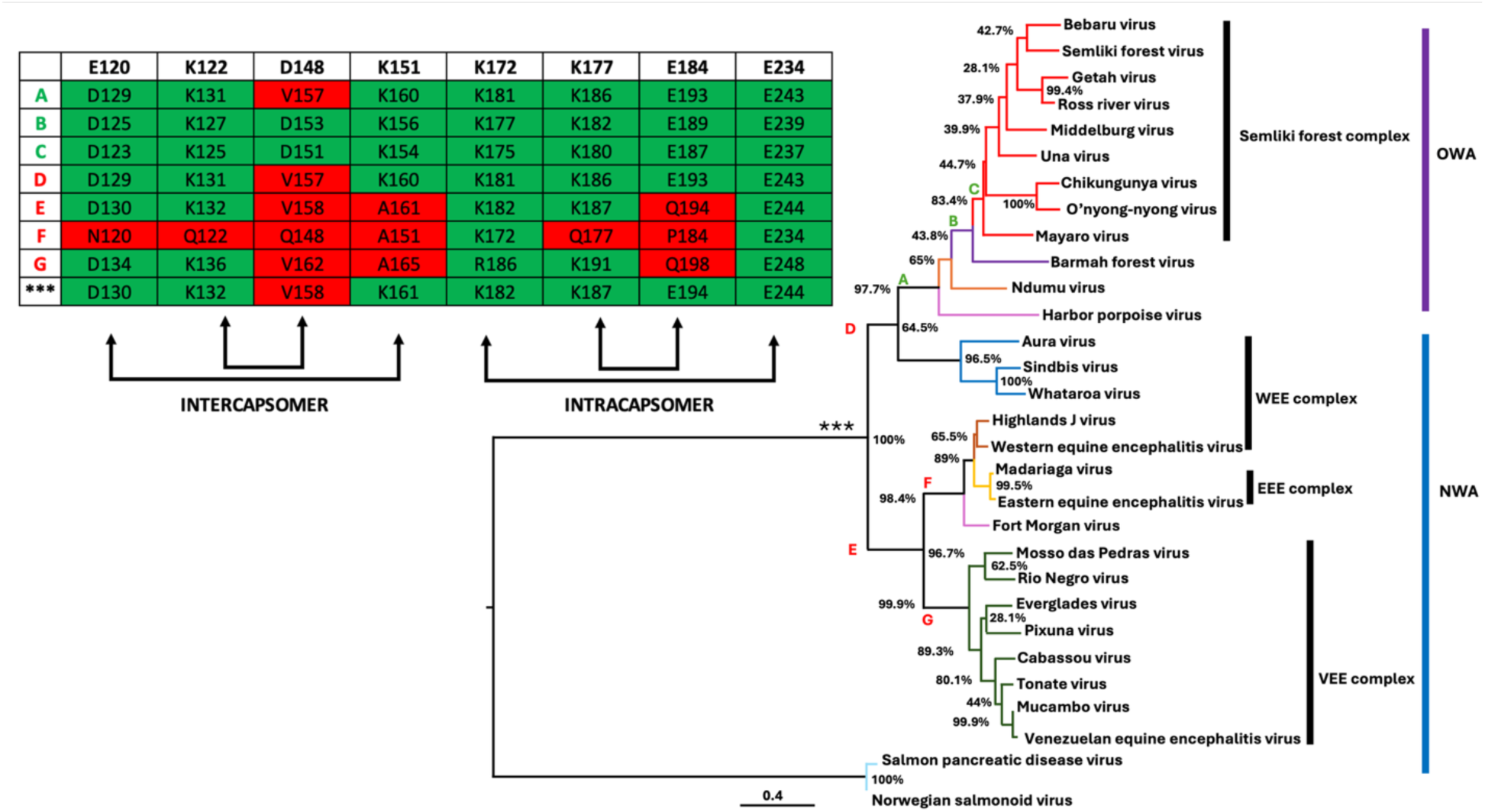
Phylogenetic tree for the Alphaviruses in Tables 1-4. The predictions of possible common ancestors made from the data in Tables 1-4 are supported by this tree. The letters refer to the nodes of interest: *** is the common ancestor of the CP for Old- and New-World Alphaviruses (OWA and NWA, respectively). The table shows the identity of the amino acids of interest in nodes *** and A-G; the green and red colors indicate whether the charge of these amino acids is conserved or not with respect to CHIVK, respectively. The colors of the branches indicate the diSerent clades. The prediction of the sequence of this node suggests that the seven amino acids of interest, which are highly conserved in Old-World Alphaviruses, were present in this ancestor; however, the origin of the amino acids observed in Old-World Alphaviruses today arose early in their evolution (see B). The node that gave origin to Sindbis is shared with Old-World Alphaviruses. At the same time, the early beginning of the rest of the New-World Alphaviruses started with a fast diversification of the intra- and intracapsomeric interactions. As these viruses evolved, the diversity of these interactions increased significantly.

Analysis of the CP results reveals that within the WEE antigenic complex, Aura, SINV, and Whataroa viruses belong to a different clade than the Highlands J and WEE viruses. Interestingly, this analysis suggests that the evolution of the CP of Fort Morgan virus differs significantly from that observed for other structural proteins. New-World Alphaviruses appear to have given rise to three main clades of the viral core. One clade includes Aura, SINV, and Whataroa viruses, which appear to be closer ancestors of Old-World Alphaviruses than the rest of the New-World Alphaviruses. This is consistent with SINV and Whataroa being found in the Old World. The second clade includes Fort Morgan, Madariaga, Highlands J, EEEV, and WEE viruses, while the third clade encompasses the rest.

Additionally, we performed an entropic and selective pressure analysis of Old- and New-World Alphavirus CPs to understand how variable these amino acids are and the types of selective pressure they may be subject to during evolution. When these sequences were analyzed, we did not detect strong selective pressure to maintain the identities of CP E120, K122, D148, K151, K172, K177, E184, and E234 (supplementary files). However, when this analysis was conducted with only Old-World Alphavirus CP, 31 sites within the CP MSA were under purifying selection, as detected by FEL or FUBAR analyses (supplementary files). Interestingly, 16 amino acids also exhibited low entropy values, indicating high evolutionary conservation. A Shannon entropy value less than 0.9 indicates high conservation; the maximum possible Shannon entropy if there are 20 possible amino acids with the same probability to appear is 4.32 (*i.e.,* random distribution), and a value of 0 refers to complete conservation. Therefore, a value of 0.9 indicates that there are only two or less possible amino acids for that position. These conserved residues are likely to be functionally or structurally important, potentially contributing to the proper functioning of these proteins. The overlap between strong negative selective pressure and low sequence variability supports the idea that these sites are subject to evolutionary constraints due to their essential roles in the viral cycle.

Notably, a general trend of low entropy was observed starting at position 130 of the 295-amino-acid alignment (of the Old-World Alphavirus consensus sequence), suggesting a region of sustained conservation across multiple species, potentially associated with a structurally constrained domain (the first 100 to 130 amino acids are disordered and interact with the genomic RNA). Furthermore, most of the sites subjected to purifying selection were located between positions 96-145 and 187-206 (supplementary files), highlighting two discrete regions of potential functional importance that may be critical for the structural integrity of the viral capsid.

A closer look at CHIKV CP in positions E120, K122, D148, K151, K172, K177, E184, and E234 showed that the Shannon entropy of these amino acids is 0.7, 0.3, 0.3, 0.3, 0.3, 0, and 0.3, respectively (see Figure 12). These results are consistent with the hypothesis that these eight amino acids are key in Old-World Alphavirus CP. The consequences of these observations will be discussed later, but they may be related to the pathogenesis and type of disease they cause.

**Figure 12.**
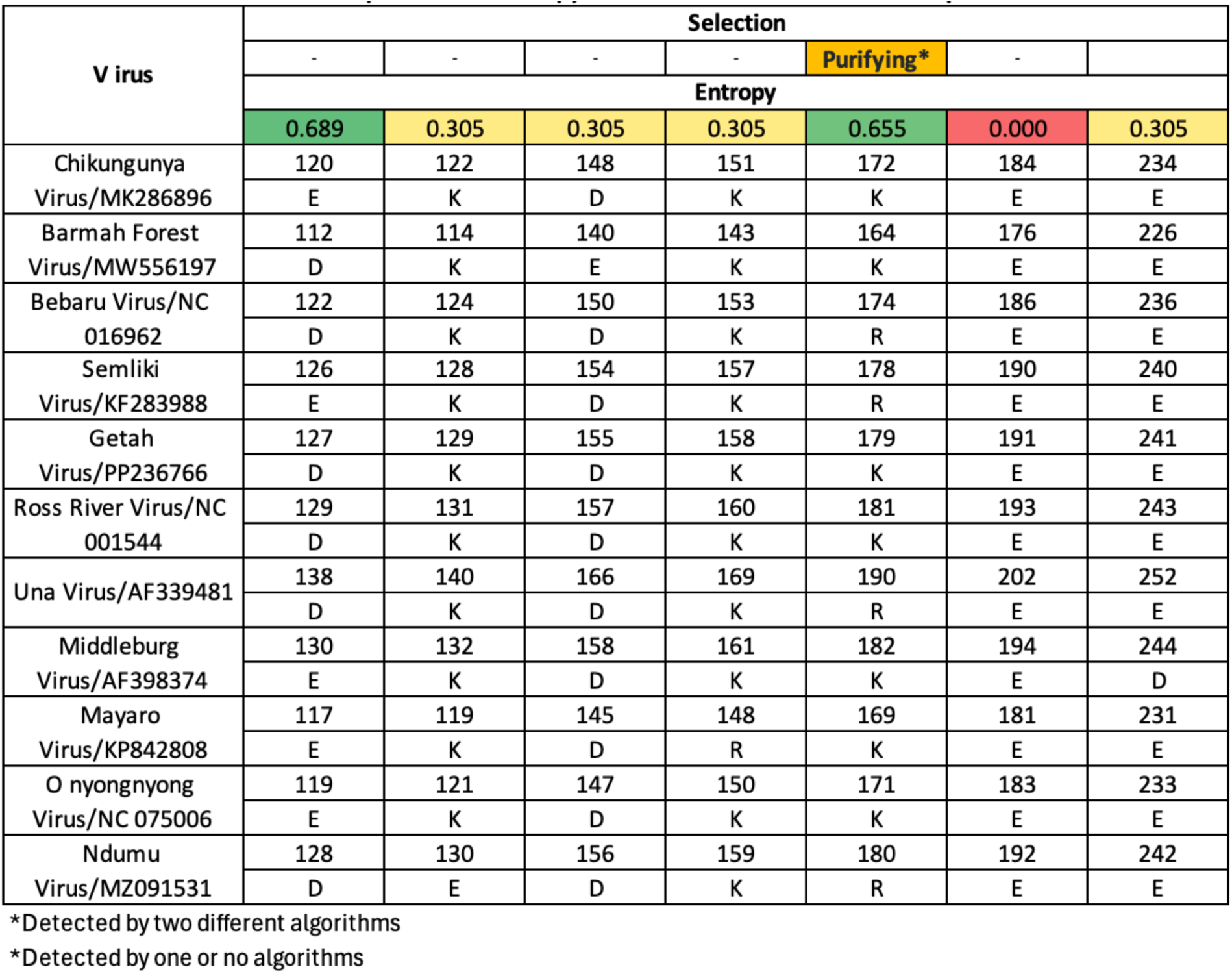
Comparison of the Shannon entropy and selective pressure of amino acids in equivalent positions to CHIKV CP E120, K122, D148, K151, K177, E184, and E234 Old-World Alphaviruses. This analysis was performed using SLAC, FEL, FUBAR, and MEME methods; sites were classified as under pervasive or episodic positive selection if detected by MEME (*p*<0.1), FUBAR (posterior probability >0.9), FEL (*p*<0.1), or SLAC (*p*<0.1).

Then we used the ExtAncSeqMEGA to infer a few most probable ancestor sequences (see the three asterisks and letters A-G in the insert in Figure 11). This analysis strongly suggests that the common ancestor of Alphaviruses that infect mammals but not fish had seven of the eight amino acids we propose mediate CP-CP and capsomer-capsomer interactions in Old-World viruses. This data implies that, after the appearance of the Old-World Alphavirus ancestor in node C, selective pressure on the eight amino acids of interest became so strong that they became invariant. The only difference between this common ancestor’s amino acids of interest and Old-World Alphaviruses is that the equivalent amino acid to CHIKV D148 is V158.

The event that led to the divergence of these two main groups resulted in a significant diversification of the New-World Alphavirus CP. However, on the one hand, the common ancestor of the Sindbis clade has the same eight amino acids as the common ancestor for all Alphaviruses that do not infect fish (*** in Figure 11). On the other hand, the common ancestor for the VEE and EEE complexes already had three mutations in the CP: one mutation in the inter- and two mutations in the intracapsomeric interactions, respectively. The most divergent node is the one that gave rise to the EEE complex.

## Discussion

The results presented here suggest that the CP-CP interactions that mediate core assembly of New- and Old-World Alphaviruses are distinct. On the one hand, based on the analysis of cryo-EM structures and the predicted structure of the CPs, it is likely that the core of CHIKV and all other Old-World Alphaviruses is controlled through highly conserved electrostatic interactions. Most likely, salt bridges control the assembly of capsomers (*i.e.,* pentamers and hexamers), and dipole-dipole-like interactions at the i3 and q3 axes are responsible for capsomer-capsomer interactions. On the other hand, the amino acids that control the assembly of capsomers and capsomer-capsomer interactions in New-World and other Alphaviruses are not conserved, and their interactions should be weaker than in the Old-World cores. Based on the proposed amino acids, it is likely that the assembly of the core of Old-World Alphaviruses is mediated by highly conserved, strong electrostatic interactions unique to this group of viruses.

Kaelber *et al*. showed that CHIKV, EEEV, and VEEV have T = 4 virions, with a minority population of T = 3 and abnormal particles (30). However, the prevalence of T = 3 and abnormal virions is higher in VEEV and EEEV than in CHIKV. While they did not propose why EEEV and VEEV virions have a higher population of misassembled virions than CHIKV, our results are consistent with a scenario where the assembly of EEEV and VEEV virions results in a smaller population of (correct) T = 4 virions than CHIKV, because the intra- and intercapsomere interactions of EEEV and VEEV CP are weaker than those of CHIKV and hence are more prone to making mistakes during assembly. Unpublished results from our group show that point mutations in the CHIKV CP at any of these eight amino acids proposed here to control the intra- and intercapsomere interactions increase the percentage of abnormal virions. While the biological implications of Kaelber and co-workers’ observations are unclear, the fact that EEEV and VEEV have a higher rate of misassembled virions could be interpreted as an attenuation mechanism. It is essential to note that EEEV and VEEV are lethal viruses, whereas CHIKV is rarely deadly (31). Therefore, there is the possibility that encephalitic Alphaviruses may be more likely to make more mistakes during assembly to achieve attenuation, thereby increasing the probability of infecting new hosts before the original host dies. However, this hypothesis is yet to be tested.

Mukhopadhyay and coworkers cryo-EM reconstructed Ross River virus and WEEV *in vitro* assembled core-like particles (CLPs) and could not achieve a resolution higher than 30 Å (7). They also attempted to reconstruct cores produced in mammalian cells, in which the lipid bilayer and the E1/E2 layer were removed, obtaining similar results to those with the CLPs. Therefore, they proposed that their inability to achieve the same resolution as intact virions suggested that CLPs deviate from an icosahedron and/or are heterogeneous (7). Recently, Chmielewski and co-workers used cryo-ET to reconstruct CHIKV particles as they budded from cultured cells (4). In this work, they demonstrated that the cores in the cytoplasm are not icosahedral, and the double icosahedral symmetry of the virion is assembled during budding. Although Chmielewski and coworkers did not investigate the molecular assembly mechanism of CHIKV virions, their results can be correlated with the hypothesis presented here. The transition from a disordered to an icosahedral core can be viewed as an annealing process in which the cytoplasmic core is not in its lowest free-energy state. It is upon interaction with the glycoproteins at the plasma membrane that multiple protein-protein interactions overcome the loss of entropy (because of the transition from disordered to icosahedral core). Based on data from Chmielewski and co-workers (4), it is reasonable to propose that weak interactions would facilitate this mechanism; for example, the proposed dipole-dipole-like interactions between capsomers depend on the distance and angle between them. An angle dependency would facilitate the reordering of a heterogeneous core into an icosahedral structure as observed by Cryo-ET (4).

We investigated the evolution of the CP, and by only focusing on the CP, we could infer that the common ancestor of non-fish-infecting Alphaviruses had almost the same putative CP-CP and capsomer-capsomer interactions as Old-World Alphaviruses. On the one hand, New-World Alphaviruses are mostly restricted to the American continent, yet there is significant variability in these interactions. On the other hand, Old-World Alphaviruses can be found in the rest of the world, and these interactions are highly conserved.

Our analysis of Shannon entropy and selective pressure clearly shows that the CPs of these two groups of Alphaviruses differ significantly; the amino acids proposed to control core assembly are invariant in Old-World viruses. Altogether, our results, along with those obtained by studying the evolution of Alphaviruses using the glycoproteins (which are responsible for viral tropism) and the virulence factor TF (17, 18), which resulted in the generation of arthrogenic and encephalitic Alphaviruses, could suggest a relationship between the CP-CP that control core assembly and pathogenicity. The event that resulted in a rapid evolution of the CP on the American continent, and allowed for weak CP-CP interactions and possibly core defects (as observed in VEEV and WEEV (30)) could have resulted in a viral attenuation mechanism. This hypothesis is consistent with the fact that the glycosylation profiles of E2 and E1 differ between these two groups of Alphaviruses. Therefore, it is possible that glycosylation changes not only affected tropism but also contributed to stabilizing cores with weak interactions (unpublished data). In other words, it is possible that the “molecular handshake” proposed by Ribeiro-Filho for MAYV (12) had to be removed to allow for assembly of the core when the interactions became weaker, and that this event might have happened when the divergence of Old- and New-World viruses occurred.

It has been proposed that Alphaviruses have a marine origin (18). However, a phylogenetic analysis of the CP alone suggests that the marine alphaviruses infecting mammals evolved later than those that infect fish. Our analysis also indicates that Harbor porpoise virus was reintroduced to marine animals after the divergence of Old- and New-World viruses. We are unclear when the Southern elephant seal virus (SEAV) was introduced. However, given that SEAV CP has six of the eight amino acids of interest conserved with CHIKV, this suggests it may have been reintroduced into a marine environment at the same time as Harbor porpoise virus.

The inference of the CP common ancestor suggests it had seven of the eight interactions we propose to control CHIKV assembly. This indicates that the event that separated Old- and New-World Alphaviruses resulted in homogeneous, and most likely more stable, Old-World cores. In contrast, this event favored increased heterogeneity in New-World Alphaviruses. The increase in variability in the intra- and intercapsomere interactions could likely be linked to the development of a lethal encephalitic disease.

## Conclusions

The interactions that control core assembly in Old-World Alphaviruses differ significantly from those of other Alphaviruses. The amino acids studied here are likely to play a crucial role in the assembly of capsomers and the core in the cytoplasm, so that they favor the transition from an irregular to icosahedral core during assembly. In Old-World Alphaviruses, these interactions are highly conserved and likely emerged early in the evolution of these viruses. The divergence of Old- and New-World Alphaviruses resulted not only in differences in tropism and disease but in structural differences. These differences could have resulted in a novel attenuation mechanism in which lethal Alphaviruses are more prone to make mistakes during assembly than non-lethal Alphaviruses. However, the results presented here require experimental confirmation, primarily because the cryo-EM data were not fully resolved and because the AlphaFold2 and 3 models were generated from the monomeric capsid protein; therefore, we may be missing protein surfaces that appear once the core is assembled. Nonetheless, combining structural prediction and MSA analyses proved to be crucial as we were able to find that only 12 out of 293 amino acids (4.1%) showed discordance in position. This suggests that combining both approaches is likely to yield a highly accurate prediction of these interactions of interest. Our group is currently testing these scenarios to investigate the contribution of these amino acids by performing site-directed mutagenesis in our CHIKV system (32) with a reporter gene to validate so we can further validate these hypothesis on core assembly, as well as the correlation between the E2 N263 glycosylation site characteristic of Old-World Alphaviruses and the identity of the amino acids that control core assembly.

## Acknowledgments

We also thank Jose Alberto Campillo-Balderas for his helpful comments during the writing of the manuscript.

