## Supplementary figures for "Highly Conserved Core Residues Define Old-World Alphaviruses and Trace Early Evolutionary Divergence"

**Supplementary Figure legends**

**Figure 1**. Comparison of the cryo-EM structure of the cores at the q3 symmetry axis of CHIKV (PDB ID: 3I2W), MAYV (7KO8), and GETV (7FD2). B) Comparison of the cores EEEV (6XO4), VEEV (7SFV), and WEEV (8DEC). Each structure has four different colors, representing that in a T = 4 icosahedron, the capsid protein (CP) in the asymmetric unit has four different chemical environments. At this symmetry site, there are large spaces between the two hexamers and the pentamer; for the Old-World Alphaviruses CHIKV, MAYV, and GETV, this space has a diameter ranging from 19 to 25 Å. For the New-World Alphaviruses EEEV, VEEV, and WEEV, the space between these three capsomers is between 13 and 22 Å.

**Figure 2.** Multiple sequence alignment of the representative consensus sequences for all the studied Alphaviruses with respect to the first 167 amino acids of CHIKV. This alignment agrees with the data obtained by using structural comparative analysis and shows how conserved the amino acids equivalent to CHIKV CP E120, K122, D148, and K151. This alignment was used to generate the color code in tables 2, 3, and 44.

**Figure 3.** Multiple sequence alignment of the representative consensus sequences for all the studied Alphaviruses with respect to amino acids 167 to 251 of CHIKV. This alignment agrees with the data obtained by using structural comparative analysis and shows how conserved the amino acids equivalent to CHIKV CP K172, K177, E184, and E234. This alignment was used to generate the color code in tables 2, 3, and 44.

**Supplementary Figure 1**

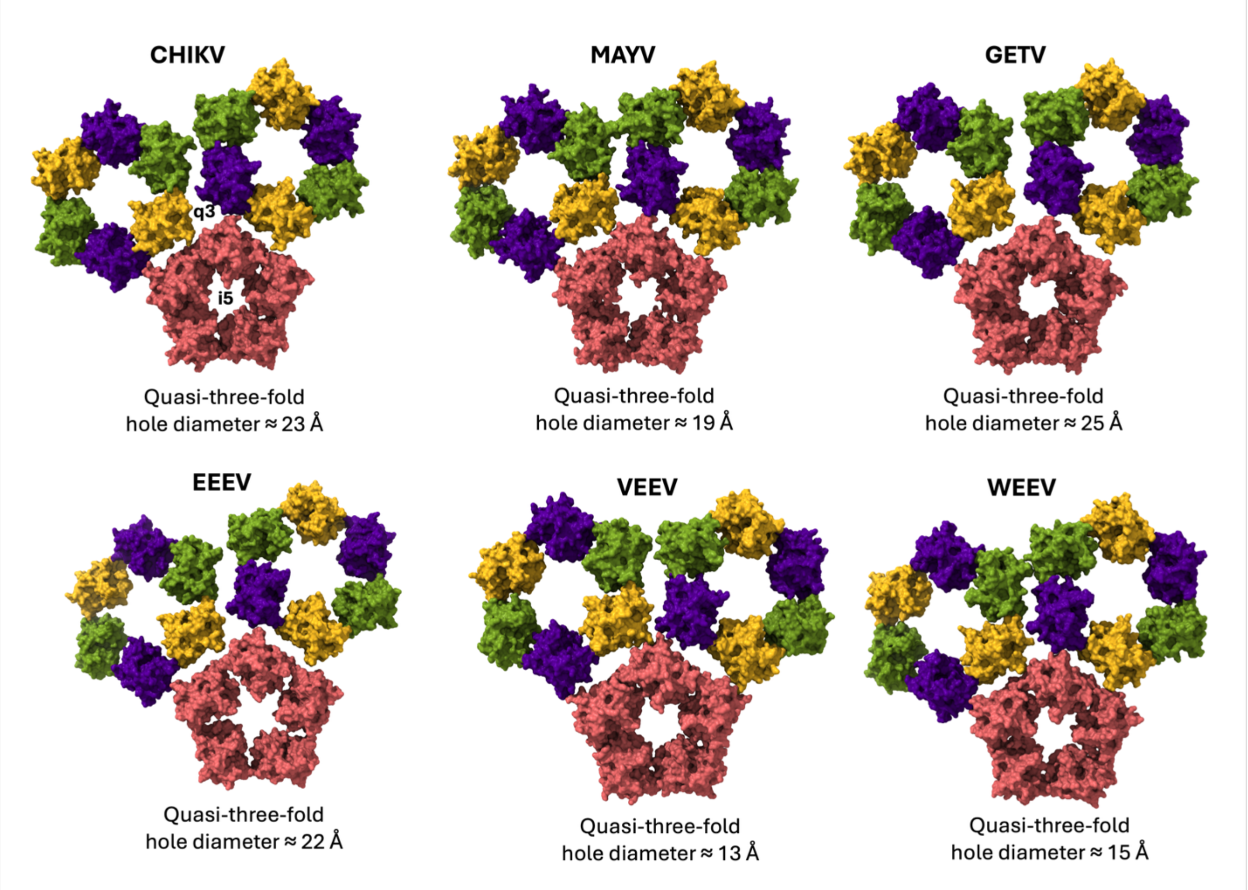

### Supplementary Figure 2

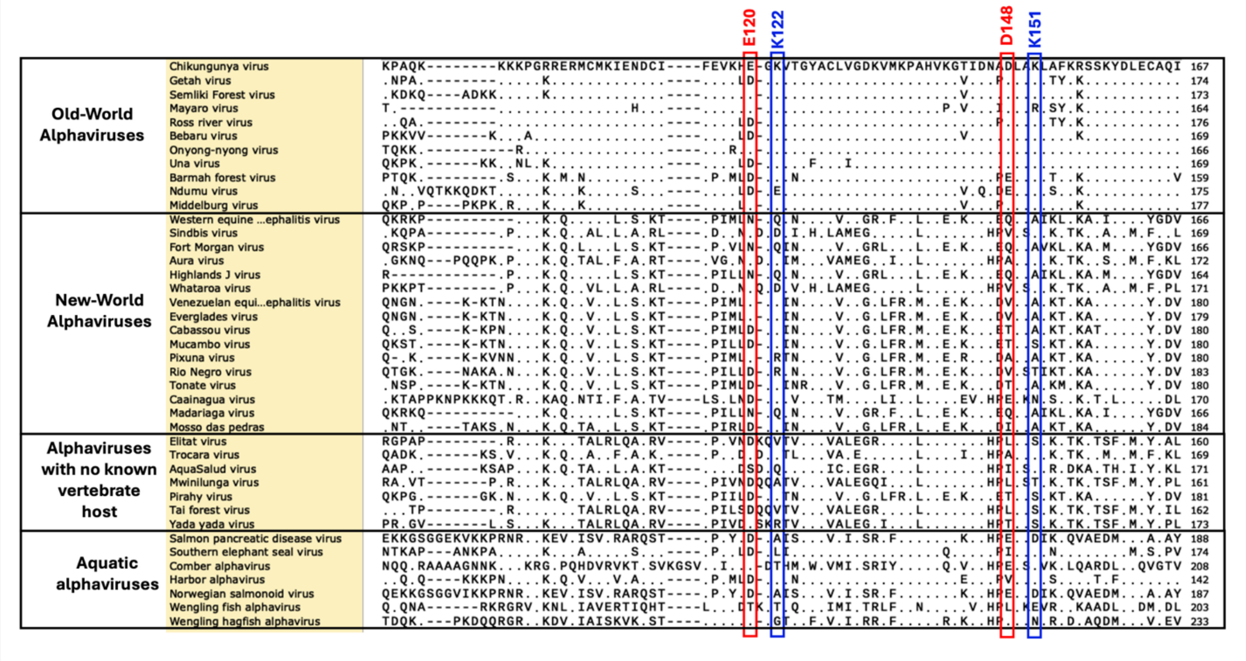

### Supplementary Figure 3

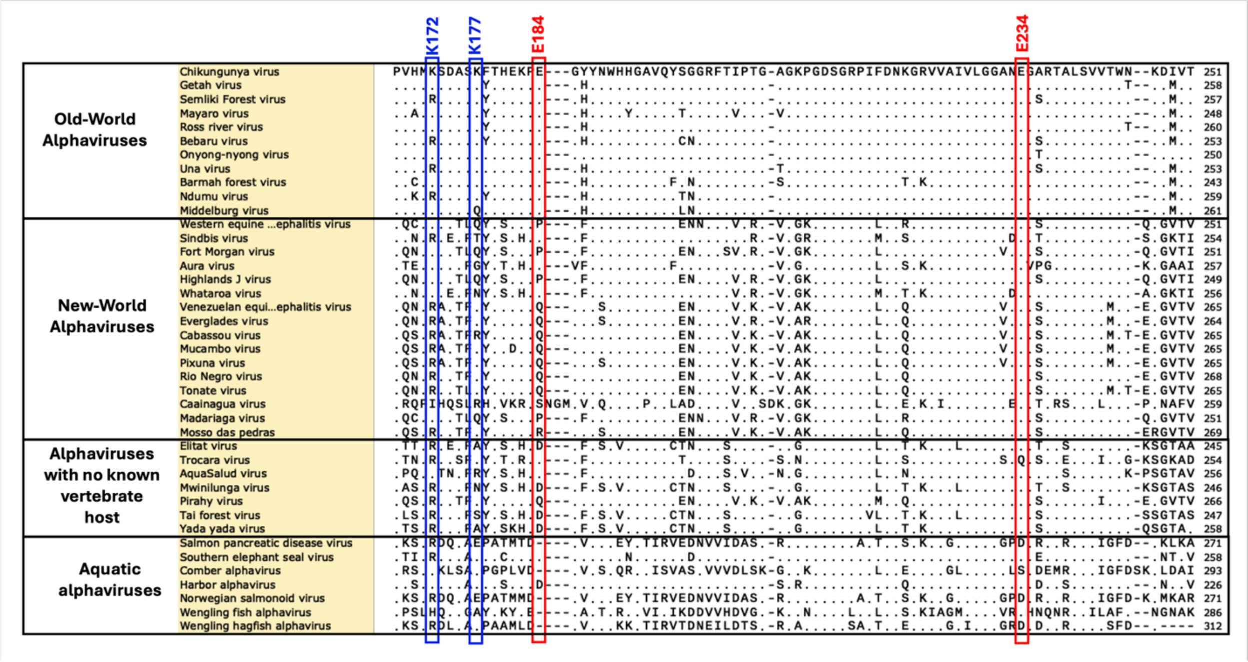
